# Kaposi’s Sarcoma-Associated Herpesvirus (KSHV) LANA Prevents KSHV Episomes from Degradation

**DOI:** 10.1101/2023.08.18.553898

**Authors:** Ken-ichi Nakajima, Tomoki Inagaki, Jonna Magdallene Espera, Yoshihiro Izumiya

## Abstract

Protein knock-down with an inducible degradation system is a powerful tool to study proteins of interest in living cells. Here, we adopted the auxin-inducible degron (AID) approach to detail Kaposi’s Sarcoma-associated herpesvirus (KSHV) latency-associated nuclear antigen (LANA) function in latency maintenance and inducible viral lytic gene expression. We fused the mini-AID (mAID) tag at the LANA N-terminus with KSHV BAC16 recombination, and iSLK cells were stably infected with the recombinant KSHV encoding mAID-LANA. Incubation with 5-phenyl-indole-3-acetic acid (5-Ph-IAA), a derivative of natural auxin, rapidly degraded LANA within 1.5 h. In contrast to our hypothesis, depletion of LANA did not trigger lytic reactivation but rather decreased inducible lytic gene expression when we stimulated reactivation with a combination of ORF50 protein expression and sodium butyrate treatment. Decreased overall lytic gene induction seemed to associate with a rapid loss of KSHV genomes in the absence of LANA. The rapid loss of viral genomic DNA was blocked by treatment with lysosomal inhibitor chloroquine. Furthermore, siRNA-mediated knockdown of cellular innate immune proteins, cyclic AMP-GMP synthase (cGAS) and Stimulator of Interferon Genes (STING), and other autophagy-related genes rescued the degradation of viral genomic DNA upon LANA depletion. These results suggest that LANA is actively protecting viral genomic DNA from sensing by cGAS-STING signaling axis, and add novel insights into the role of LANA in latency maintenance.

**IMPORTANCE:** KSHV LANA plays a wide variety of roles in latency maintenance and lytic gene expression. We adopted the inducible protein knockdown approach and revealed that depletion of LANA induced rapid degradation of viral genomic DNA. The viral genome degradation was rescued by inhibition of the cellular innate immune pathway and autophagy. These observations suggest that LANA might play a role in hiding KSHV episome from cellular innate immune DNA sensors. Our study thus provides novel insights into the role of LANA in latency maintenance.

## INTRODUCTION

Kaposi’s sarcoma-associated herpesvirus (KSHV) is an etiological agent of Kaposi’s sarcoma (KS), primary effusion lymphoma (PEL), multicentric Castleman’s disease (MCD), and KSHV inflammatory cytokine syndrome (KICS) (1). KSHV is one of a few pathogens recognized as a direct carcinogen, which also includes hepatitis B virus (HBV), human papillomavirus (HPV), and Epstein-Barr virus (EBV) (2–6). Significant effort has been taken to help patients by finding a cure for devastating diseases, but we have not been successful yet. Achieving a better understanding of essential viral proteins for the KSHV life cycle is therefore critically important.

Like other herpesviruses, KSHV exhibits a biphasic life cycle consisting of a life-long latent infection phase and a transient lytic reactivation phase, which are distinguished by their gene expression profiles (7–10). During the latent phase, the KSHV genomic DNA persists as a circular episome in the host cell’s nucleus (11–14), and the majority of viral gene expression is silenced (14–18). Upon reactivation from latency, the full repertoire of inducible viral genes is activated in a temporally-regulated manner, leading to the transcriptional activation of three classes of lytic genes, referred to as immediate early, early, and late genes (8, 10). ORF73 is one of the few genes expressed in the latent phase and encodes the latency-associated nuclear antigen (LANA). Previous studies have shown that LANA is necessary for the persistence of the viral genome in an episomal state. LANA directly binds to the conserved terminal repeat (TR) sequences in the KSHV genome through its C-terminal domain and docks onto the host chromosome through its N-terminal chromatin binding domain, thus allowing the viral genome to hitch a ride on the host chromosome during mitosis and maintain stable episomal copy numbers in the latently infected cells (19, 20). The presence of LANA is sufficient to mediate the replication and maintenance of a plasmid containing the KSHV latent origin of replication (ori-P) in transfected cells (21, 22). LANA tethers KSHV epsiomes by interacting with cellular histone H2A and H2B, and also recruits cellular DNA replication machinery to TRs during the S-phase of the cell cycle (23). LANA also forms a complex with many histone enzymes such as bromodomain-containing proteins 2 and 4 (BRD2/4), KDM3A, Polycomb repressor complex 2, hSET1 complex, and MLL1 complex (24–31). It has been proposed that LANA antagonizes the transcription of lytic genes during latency, and facilitates the establishment of latent infection via multiple mechanisms. For example, LANA inhibits K-Rta expression, a viral transactivator essential for lytic reactivation, by repressing the transcriptional activity of the K-Rta promoter (32). Another mechanism includes recruitment and tethering of PRC2, CTCF, and cohesin (8, 33–35). These studies primarily utilized gene knockdown or knock-out that may have introduced indirect effects to compensate for the loss of LANA biological activity during the establishment of the cell clones.

RNA interference (RNAi)-mediated gene silencing and CRISPR/Cas-mediated gene knockout have been utilized widely to study the function of a specific gene/protein of interest. However, some genes that are essential for cell growth or survival are difficult to be silenced transcriptionally or knocked out. In contrast, inducible protein degradation, also known as protein knockdown, is a versatile approach for studying the function of a specific gene/protein that is essential for cell growth (36). The auxin-inducible degron (AID) system has recently emerged as a powerful tool to conditionally deplete a target protein in various organisms, and the AID-tagged target protein can be degraded within a few minutes to a few hours after the addition of the plant hormone auxin (37, 38). Mechanistically, rice-derived TIR1 (OsTIR1) protein interacts with endogenous Skp1 and Clu1 proteins to form a functional Skp1-Clu1-F-box (SCF) E3 ubiquitin ligase complex in non-plant cells (see (37)). The OsTIR1-containing SCF E3 ubiquitin ligase is activated only when the plant hormone auxin (or its derivatives such as 5-phenyl-indole-3-acetic acid; 5-Ph-IAA) is present. The target protein will then be polyubiquitinated and degraded by the proteasome. Because the knockdown of proteins through inhibition of transcription would take some time, and cells often adapt to the changes by compensating for the biological effects through other means. In contrast, rapid protein depletion bypassed these indirect and undesired effects and therefore allows researchers to identify more direct biological effects of the proteins of interest. KSHV LANA is a multi-functional protein and therefore suitable for applying this approach to dissect its different biological functions from one another.

Cellular innate immunity is the first line of defense against incoming pathogens, including bacteria and viruses. The innate immune system is an evolutionally conserved host defense with key features being shared between plants and animals (39, 40). Several fundamental molecules/mechanisms for cellular innate immunity responses have been identified, including pattern recognition receptors (PRRs) signaling (e.g., Toll-like receptors (TRLs), nucleotide oligomerization domain-like receptors (NLRs), and RIG-I-like receptors) (41) and a DNA sensor cGMP-AMP synthase (cGAS) (42). PRRs recognize microbial components known as pathogen-associated molecular patterns (PAMPs) (41). Examples of these include bacterial cell wall components such as lipopolysaccharides (LPS), and ribonucleic acids (RNA) derived from viruses. PRRs recognize and bind their respective ligands and recruit adaptor molecules through their effector domains, and then initiate downstream signaling pathways to exert effects. On the other hand, cGAS has recently emerged as a non-redundant DNA sensor invaluable for detecting many pathogens (42). The binding of cGAS to double-stranded DNA (dsDNA) allosterically activates catalytic activity and leads to the production of 2’3’ cyclic GMP-AMP (2’3’-cGAMP) from ATP and GTP, which acts as a second messenger and potent agonist of the downstream protein, Stimulator of Interferon Genes (STING). For this reason, cGAS can recognize a broad range of DNA species of both foreign and self-origin. Binding to 2’3’-cGAMP triggers conformational changes in STING, followed by its trafficking from the endoplasmic reticulum (ER) to the Golgi apparatus via the ER-Golgi intermediate compartment (ERGIC) (42). Upon reaching the ERGIC/Golgi, STING is oligomerized into a signaling platform that recruits downstream proteins to initiate an immune response, including the cytokine production (42). In addition, a recent study revealed that STING also induced the autophagy pathway to degrade components derived from pathogens as a cellular defense response (43).

In this study, we constructed a novel KSHV BAC16 (bacterial artificial chromosome 16), in which the N-terminus of LANA is tagged with a 7 kDa mini-AID (mAID) tag. We assessed the specific contribution of LANA for maintaining viral episomes in infected cells and inducible lytic gene regulation during reactivation.

## RESULTS

### Generation and characterization of mAID-LANA BAC16

Protein knock-down with the degron system is an alternative approach to dissecting the function of proteins of interest in living cells. The protein knock-down is especially suitable for proteins that are difficult to establish stable knock-down or knockout from cells. LANA is an abundantly expressed protein in KSHV latently infected cells and plays an invaluable role in maintaining latency by at least two molecular mechanisms: (i) tethering KSHV DNA to host chromosomes to maintain viral genomes and (ii) suppressing inducible lytic gene expression. To reveal LANA’s contribution to latency maintenance more directly, we adapted the inducible protein degradation approach to a recombinant KSHV BAC system and established recombinant KSHV which encodes mAID-fused LANA (mAID-LANA). We first prepared a template plasmid which is used to generate PCR fragments for recombination. The template plasmid contains a kanamycin resistance cassette in the mAID coding sequence as an excisable format with I-SceI induction. The mAID-kanamycin DNA fragment was first amplified with primers with ∼100 bp homology arms, and amplified fragments were then used for recombination by using a two-step recombination method as previously described (44–46) **(Fig. 1A)**. The recombination junction and adjacent regions were amplified by PCR, and the PCR fragments were directly sequenced to confirm in-frame insertion. The mAID-LANA BAC16 was transfected into iSLK cells to generate iSLK cells harboring mAID-LANA BAC16. The cells were further transduced with a recombinant lentivirus expressing FLAG-tagged rice-derived TIR1 F74G (FLAG-OsTIR1 F74G) protein. We named this cell subline as iSLK-OsTIR1-mAID-LANA BAC16 cell. OsTIR1 is a high-affinity auxin binding protein that interacts with endogenous Skp1 and Cul1 proteins to form a functional SCF E3 ubiquitin ligase complex (37). The OsTIR1-containing SCF E3 ligase is only activated in the presence of plant hormone auxin or its derivatives and polyubiquitinates AID-tagged (or mAID-tagged) target protein which leads to rapid proteasome-mediated degradation. We next verified the virological function of mAID-LANA. We first compared the amount of KSHV virion productions from iSLK-OsTIR1-mAID-LANA BAC16 cells with that from iSLK.219 cells **(Fig. 1B).** We also examined the infectivity of progeny virions produced from iSLK-OsTIR1-mAID-LANA BAC16 cells. For that, iSLK cells were infected with purified mAID-LANA BAC16 or BAC16 wild-type virions at a multiplicity of infection (MOI) of 10, and GFP-positive iSLK cells were quantified by flow cytometry. The GFP-positive (infection) ratio was comparable with that of the BAC16 WT virus **(Fig. 1C)**. Furthermore, we confirmed that KSHV episomes were maintained during cell passage. These results suggested that the tagging with the 68-amino acid residue mAID at the N-terminus did not interfere with the LANA function.

**Fig. 1.**
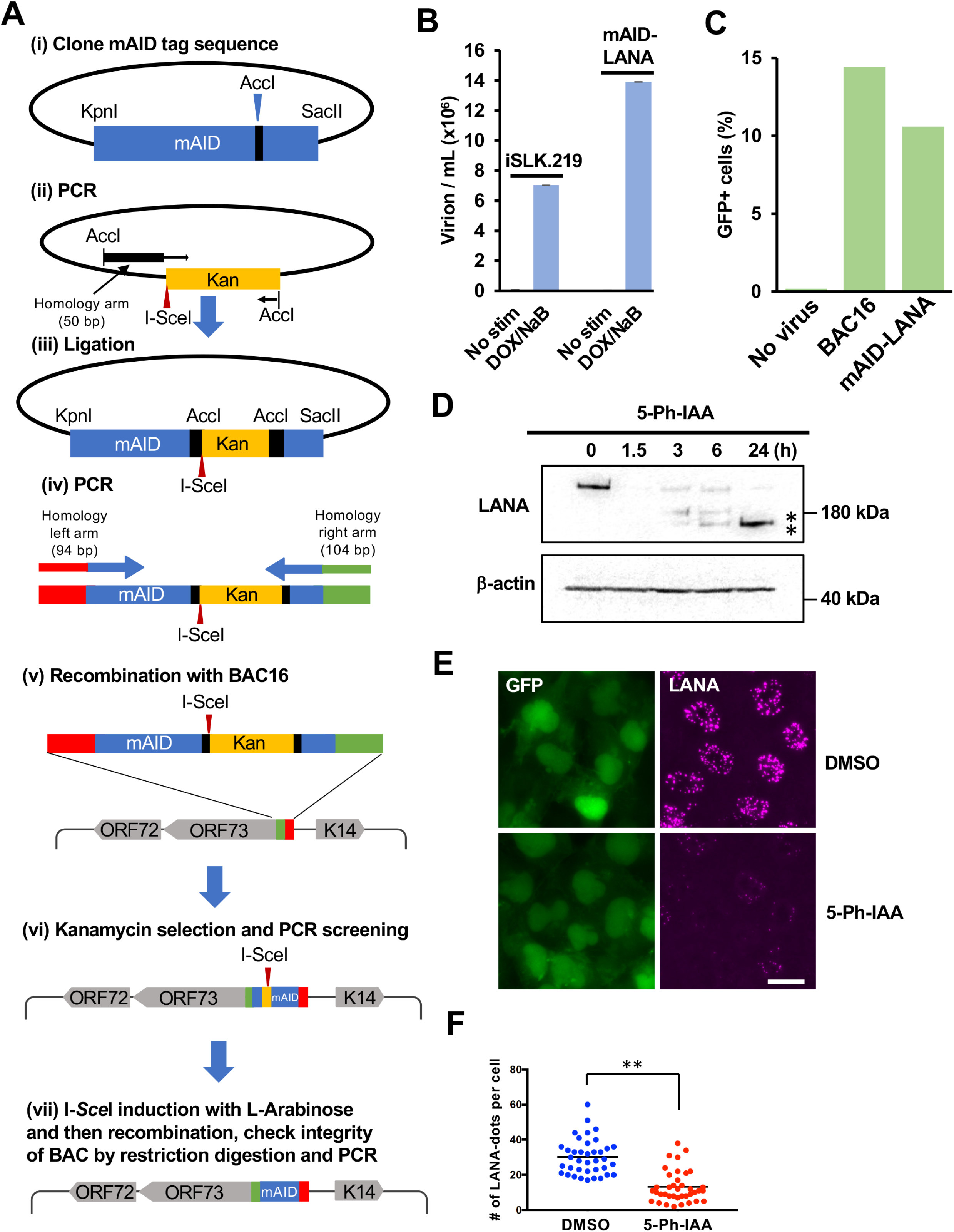
Generation of mAID-LANA BAC16. (A) The schematic diagram for construction of mAID-LANA with KSHV BAC16. (i) The codon-optimized cDNA fragment for mAID (mAID KpnI-SacII pBS fragment) was synthesized and cloned into the pBlueScript vector between the KpnI and SacII restriction enzyme sites. (ii) The kanamycin cassette with I-SceI recognition sequence along with 50 bp of the homologous sequence was generated by PCR with pEP-Kan plasmid as a template and mAID-SacII Fragment KanFw and mAID-SacII Fragment KanRv as primers, and cloned into the AccI restriction enzyme site. (iii to v) The resulting plasmid was fully sequenced and used as a template to generate a DNA fragment for homologous recombination with BAC16 inside bacteria. mAID-ORF73N-Fw and mAID-ORF73N-Rv were used as primers. (vi and vii) After confirmation of insertion at the correct site by colony PCR screening, the kanamycin cassette was deleted by recombination with induction of I-SceI in bacteria by incubation with L-arabinose. Correct insertion of the mAID-tag and integrity of BAC DNA were confirmed by sequencing of the PCR-amplified fragment and restriction digestions. Primers and the DNA fragments used are listed in Table 1. (B) Production of progeny virus. Capsidated viral DNA copy number was quantified by qPCR at 96 h post-stimulation. (C) *De novo* infection. iSLK cells were infected with the purified virus at an MOI of 10, and the GFP-positive cell population was quantified by flow cytometry 48 h after infection. (D) Depletion of mAID-LANA by 5-Ph-IAA treatment. iSLK-OsTIR1-mAID-LANA BAC16 cells were treated with 2 μM 5-Ph-IAA, and LANA depletion was assessed by Western blotting. Asterisks indicate possible degradation product(s). (E) Fluorescence micrographs of cells treated with 5-Ph-IAA (2 μM) for 24 h. LANA was visualized by immunofluorescence staining. Bar, 20 μm. (F) Quantification of LANA-dots. The number of LANA-dots per nucleus was manually counted. **, *p*<0.01.

**Table 1.**
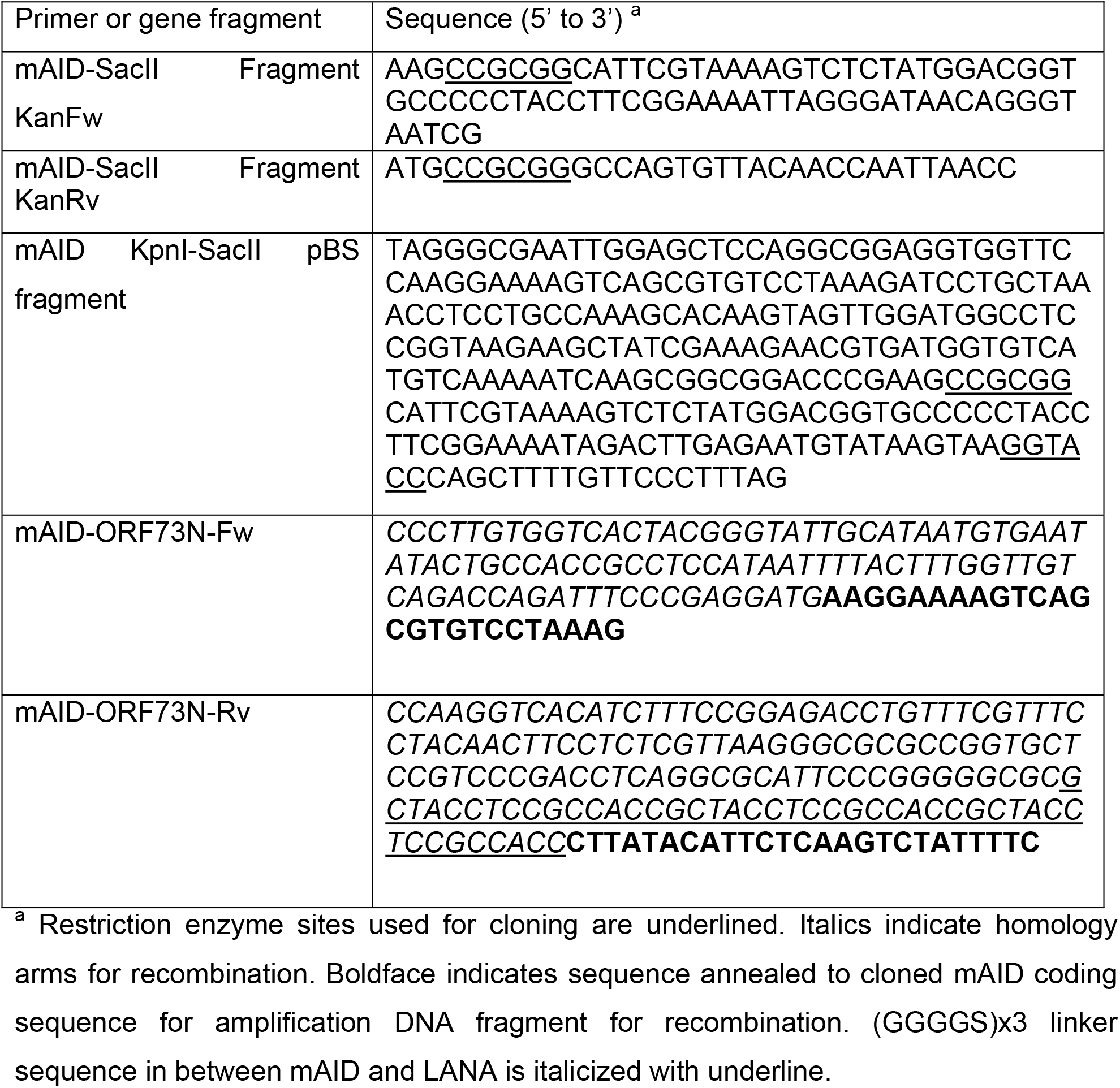

### Rapid degradation of LANA with 5-Ph-IAA

Next, we examined how quickly and to what degree incubation with auxin can deplete LANA. iSLK-OsTIR1-mAID-LANA BAC16 cells were incubated with 2 μM 5-phenyl-indole-3-acetic acid (5-Ph-IAA), a derivative of the natural auxin indole-3-acetic acid (IAA), for 0, 1.5, 3, 6, and 24 h. The cells were lysed, and an equal amount of protein was subjected to SDS-PAGE followed by Western blotting with an anti-LANA antibody. As shown in **Fig. 1D**, LANA was quickly depleted within 1.5 h of incubation with 5-Ph-IAA. LANA directly binds to the terminal repeat (TR) region of viral genomic DNA as multimers and binding to the 30-40 copies of the TR sequences concentrates LANA on the viral genome that makes them visible as “LANA-dots” or “LANA-speckles” (47) (**Fig. 1E**). Thus, a LANA-dot also indicates a single viral episome in the nucleus (21). As expected, the LANA-dots signal was drastically reduced at 24 h after the addition of 5-Ph-IAA (**Figs. 1E and 1F**). We counted the number of LANA-dots, and the result shows that the number of LANA-dots per cell was also significantly decreased by 5-Ph-IAA treatment in addition to intensity of LANA staining (**Fig. 1F**). In addition, the GFP fluorescence signal was decreased at 24 h after addition of 5-Ph-IAA (**Fig. 1E**). These results indicated that depletion of LANA might lead to elimination of viral episome in the cell.

### Viral lytic gene expression after depletion of LANA

We next examined the effects on gene transcription to assess the gene silencing function of LANA. We reactivated iSLK-OsTIR1-mAID-LANA BAC16 cells after the depletion of LANA, and the cells were fixed and stained to visualize viral lytic protein expression. The cells were first treated with or without 5-Ph-IAA for 24 h, and reactivated with doxycycline and sodium butyrate for another 24 h. Fluorescence of K8α was seen in approximately 40% of cells without depletion of LANA (the second row in **Fig. 2A**), and this observation is consistent with previous studies, in which approximately 30-40% of cells expressed early gene products such as ORF6 (45). To our surprise, the depletion of LANA itself did not globally induce lytic reactivation (the third row in **Fig. 2A**), but rather inhibited expression of K8α (the bottom row in **Fig. 2A**) when lytic reactivation was induced with doxycycline and sodium butyrate. Viral lytic gene expression was also confirmed by Western blotting (**Fig. 2B**) and real time-qPCR (**Fig. 2C**). Expression of K8α was still seen at 1.5, 3, and 6 h after the addition of 5-Ph-IAA compared to the control when cells were reactivated. Consistent with the immunofluorescence study (**Fig. 2A)**, K8α expression was substantially decreased at the 24 h time point (**Fig. 2B**). mRNA expression for lytic genes was reduced to less than one-fourth to one-tenth of control levels at 24 h in reactivated, LANA-depleted cells (**Fig. 2C**). However, two highly inducible non-coding RNAs (PAN RNA and T1.5) showed slightly increased in the absence of LANA (Fig. 2C, 5-Ph-IAA (+) Dox/NaB (-)) even though many of other lytic ORF expression were decreased. The results suggested that LANA depletion alone activated transcription of limited KSHV genomic domains.

**Fig. 2.**
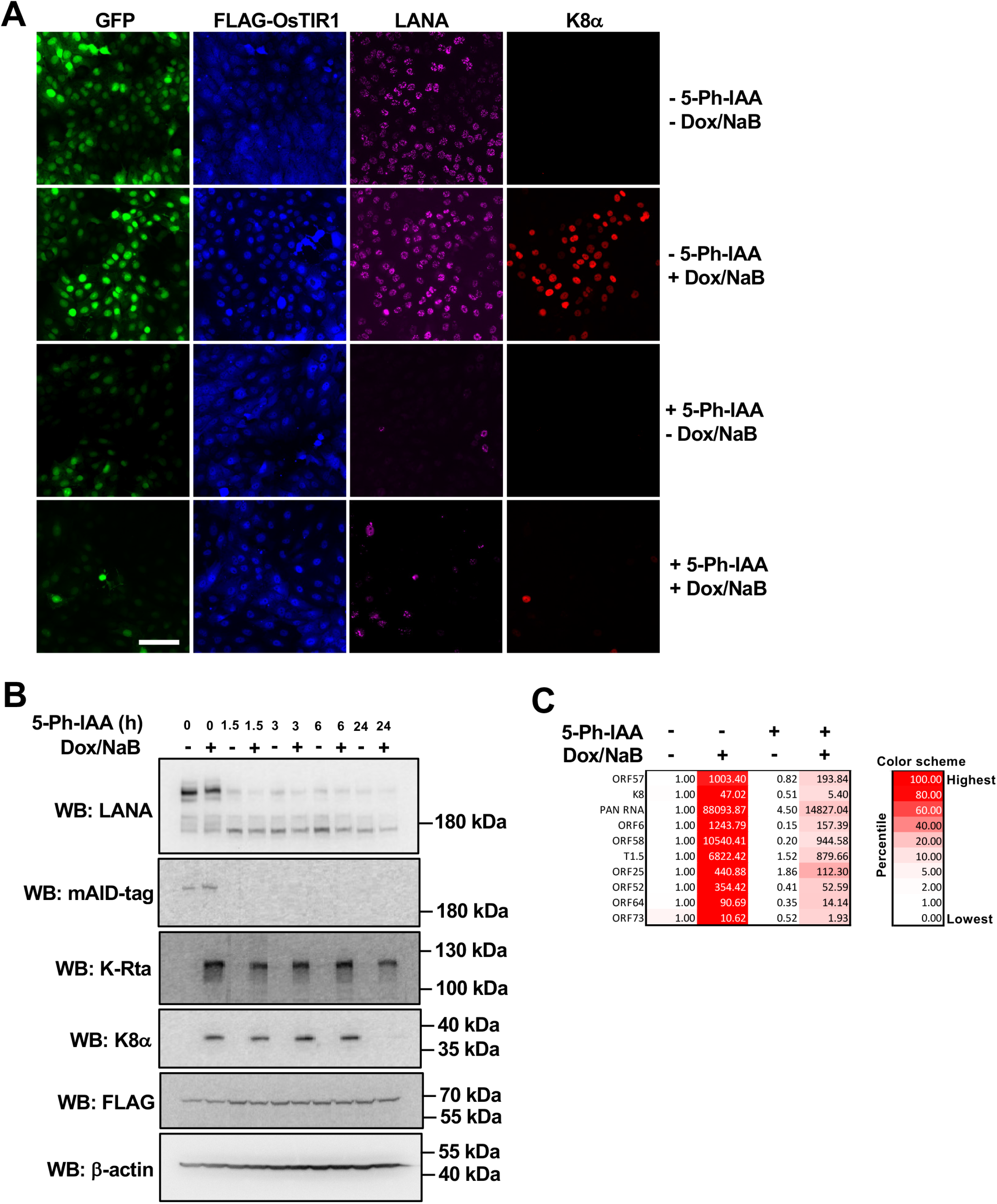
Lytic reactivation after depletion of LANA. (A) Immunofluorescence microscopy. iSLK-OsTIR1-mAID-LANA BAC16 cells were treated with or without 2 μM 5-Ph-IAA for 24 h, treated with or without 1 μg/ml doxycycline plus 1.5 mM sodium butyrate (± Dox/NaB) for another 24 h, and then subjected to immunofluorescence staining with anti-FLAG, LANA and K8α antibodies. Bar, 100 μm. (B) Western blotting. iSLK-OsTIR1-mAID-LANA BAC16 cells were treated with 2 μM 5-Ph-IAA for 0, 1.5, 3, 6, and 24 h, and then treated with or without 1 μg/ml doxycycline plus 1.5 mM sodium butyrate (± Dox/NaB) for 24 h. The total cell lysate was prepared and subjected to immunoblotting with indicated antibodies. (C) Real-time qPCR. Total RNA was extracted and subjected to real-time qPCR. 18s rRNA was used as an internal standard for normalization, and the 5-Ph-IAA (-) Dox/NaB (-) was set as 1.

### Viral episome degradation upon depletion of LANA

Treatment with 5-Ph-IAA reduced LANA-dots signal as well as GFP fluorescence signal in the cells (**Figs. 1E and 1F**), suggesting that depletion of LANA presumably decreased viral episomal copies within the cell. Accordingly, we next determined KSHV genomic copy number per cell by normalizing it to the host genome. The results showed that the depletion of LANA indeed induced a rapid reduction of viral genomic DNA to approximately 25% of control within 24 h (**Fig. 3A**). The results were somewhat surprising since iSLK cells are unlikely to divide twice within 24 h. If failure of episome tethering is the only reason for the episome loss, we speculate that having 1/4 of episomes would require at least 2 cell divisions. We therefore counted the number of cells in the same condition, and the result shows that the number of cells only doubled after 24 h (**Fig. 3B**), suggesting that these cells divided once in 24 h. We depicted a theoretical graph showing putative DNA copy number at 24 h with or without active DNA degradation, when cells divide once in 24 h (**Fig. 3C**). These observations collectively suggested that active viral episome degradation might be occurring in the absence of LANA in the cells.

**Fig. 3.**
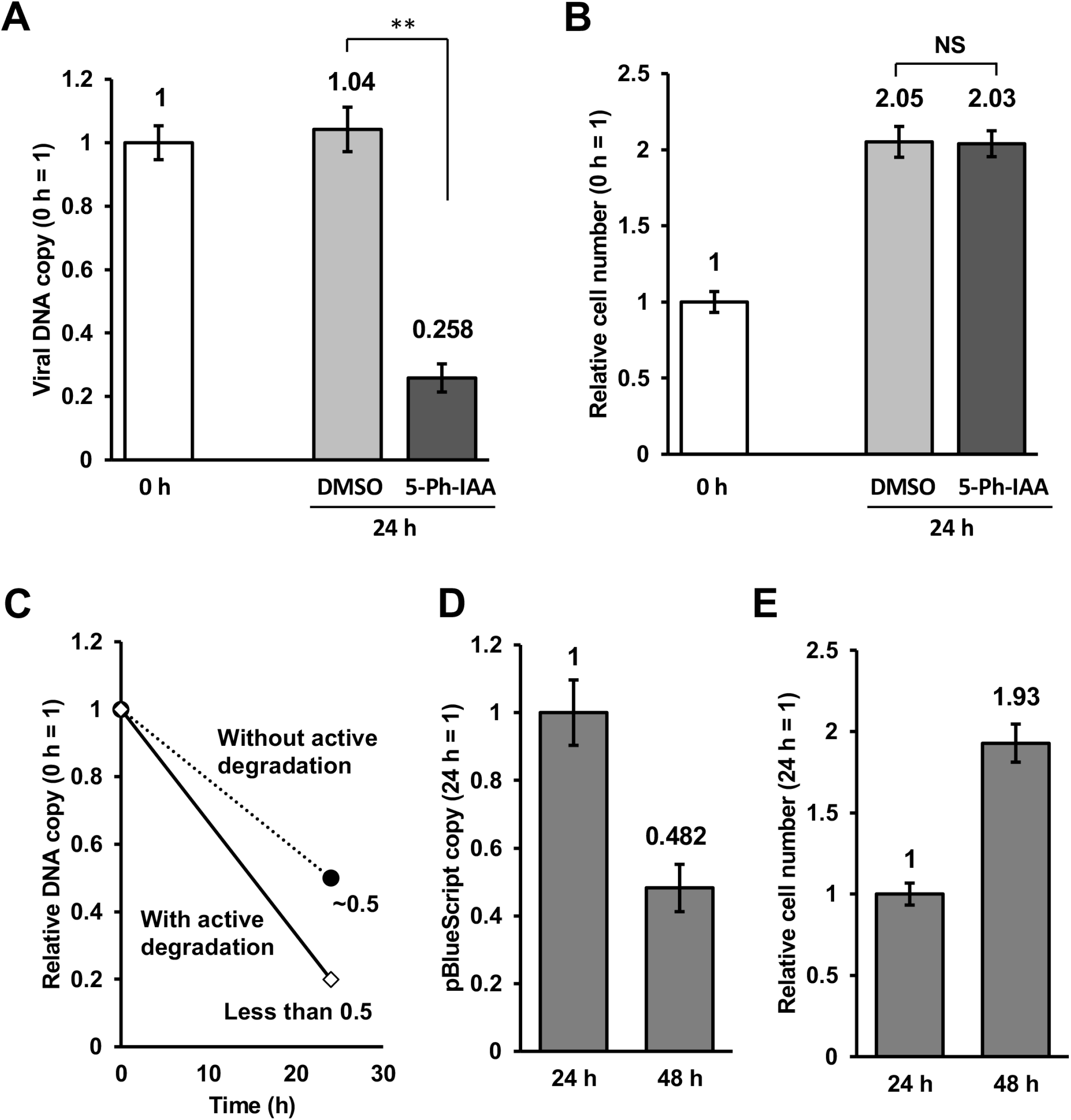
(A and B) Rapid reduction of viral DNA copy after depletion of LANA. iSLK-OsTIR1-mAID-LANA BAC16 cells were treated with DMSO (0.1%) or 2 μM of 5-Ph-IAA for 24 h, and then followed by the indicated analyses. (A) Total DNA was extracted, and viral DNA copy was measured by real-time qPCR. 18S rRNA was used as an internal standard for normalization. **, *p*<0.01. (B) Cells were trypsinized and suspended in 1% BSA, 0.1 mM EDTA in PBS. The number of cells was manually counted by using a hemocytometer. The number of cells at 0 h was set as 1. NS, not significant. (C and D) Copy number of pBlueScript vector in iSLK-OsTIR1-mAID-LANA BAC16 cells. The cells were transfected with 2 μg of pBlueScript KS+ with Lipofectamine 2000 reagent, incubated for 24 h and 48 h, and followed by the indicated analyses. (C) Cells were trypsinized, and treated with DNase I in order to digest DNA that was present outside of the cells. Total DNA was extracted, and a copy of pBlueScript was measured by real-time qPCR. We used a primer pair for β-lactamase gene encoded by the pBlueScript vector. 18S rRNA was used as an internal standard for normalization. (D) Cells were trypsinized and suspended in 1% BSA, 0.1 mM EDTA in PBS. The number of cells was manually counted by using a hemocytometer. The number of cells at 24 h was set as 1.

We next asked if the DNA degradation was specific to the viral episome. We transfected pBlueScript KS+ plasmid into iSLK-OsTIR1-mAID-LANA BAC16 cells, extract DNA from the transfected cells at 24 h and 48 h after transfection, and quantified the copy of pBlueScript plasmid. If the transfected pBlueScript was actively degraded in the cells, we expect a similar decline in copy number. We measured the DNA copy at two-time points (24 h and 48 h after transfection). As shown in **Fig. 3D**, pBlueScript copy number at 48 h was 0.48, when we set the copy number at 24 h as 1. We also counted the cell number in the same condition, and the result showed that the relative cell number at 48 h was 1.9 when we set the cell number at 24 h as 1 (**Fig. 3E**). The results suggested that in the absence of LANA, KSHV episome is more preferentially depleted in iSLK cells.

### Viral episome degradation in the lysosome

Next, we examined how viral episomes were degraded upon LANA depletion. Lysosome is an organelle in which various endogenous molecules, as well as exogenous molecules, are degraded (48). Particularly, the lysosome plays a crucial role in autophagy-mediated degradation (49). The pH of the lysosomal lumen is kept at 4.5-5 while the cytosolic pH is approximately 7.4 (50). The luminal acidic pH is critical for hydrolases localized in the lysosomal lumen (e.g., proteases and the lysosomal DNase II) that decompose many molecules, organelles, and pathogens and their constituents (48). Chloroquine is a lysosomotropic agent that was historically developed as an anti-malarial drug (51). Chloroquine passively diffuses into the lysosome, where it accumulates as a protonated form. This results in an increase of the intra-lysosomal pH, hence the inhibition of lysosomal hydrolases that require an acidic pH for their proper function (52).

We first treated the iSLK-OsTIR1-mAID-LANA BAC16 cells with chloroquine for 30 min, and then 5-Ph-IAA was added. The cells were further incubated for 24 h in the presence of chloroquine and 5-Ph-IAA. If viral DNA is degraded in lysosome upon LANA depletion, chloroquine treatment would rescue the reduction, and the viral DNA copy should be approximately 0.5 if the cells divided once in 24 h. As shown in **Fig. 4A**, viral DNA copy was recovered to 0.477 from 0.252 by chloroquine treatment. Note that chloroquine treatment itself did not have effects on viral DNA copy number. We also counted the number of cells in the same condition, and results showed that these cells still divided once in 24 h (**Fig. 4B**). These results suggest that viral DNA was delivered to the lysosome for degradation in the absence of LANA.

**Fig. 4.**
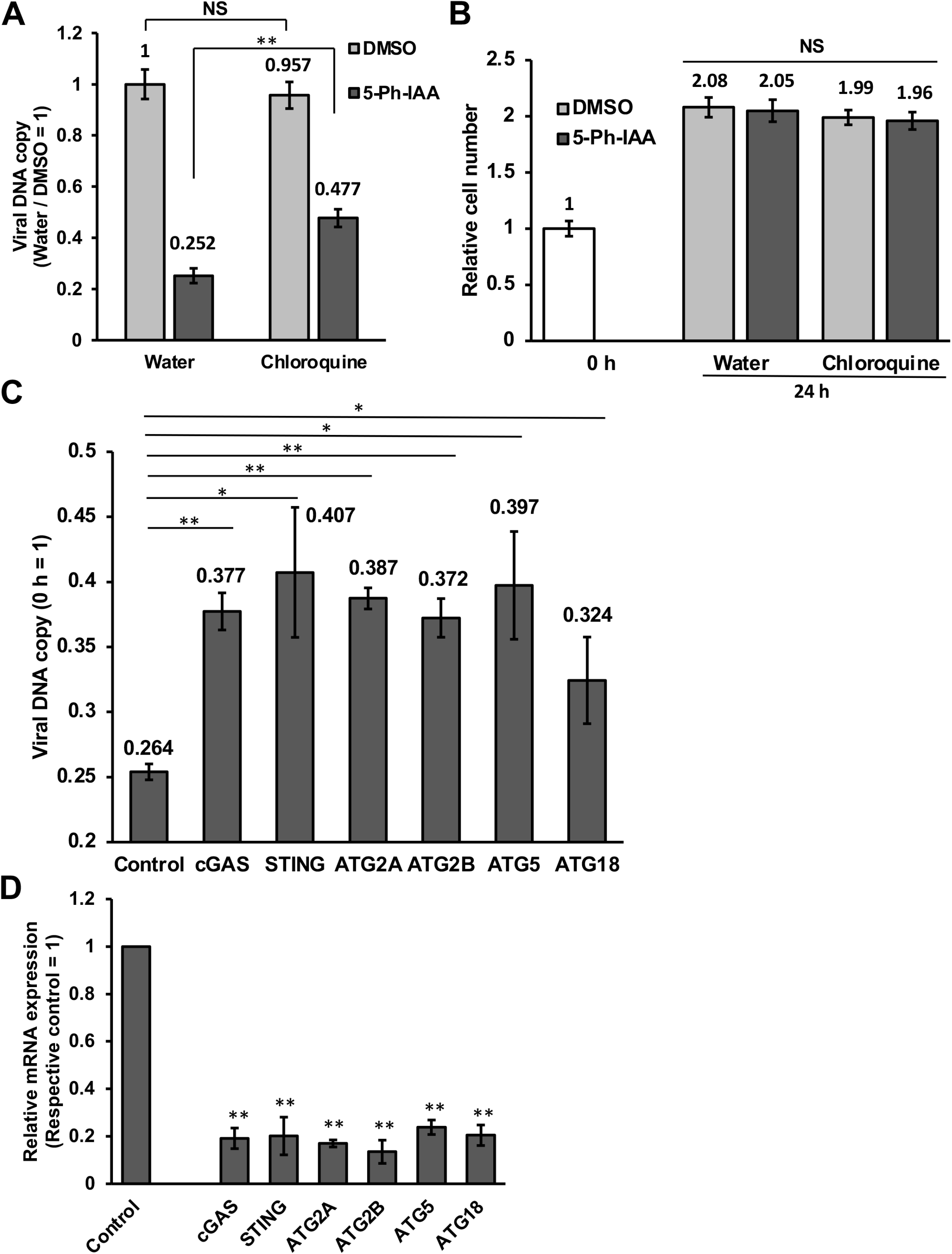
Viral DNA degradation is mediated by cGAS-STING and autophagy pathway. (A) Treatment with chloroquine rescued the reduction of viral DNA upon LANA depletion. iSLK-OsTIR1-mAID-LANA BAC16 cells were treated with 50 μM of chloroquine for 30 min, and then 5-Ph-IAA was added to a final concentration of 2 μM. At 24 h after the addition of 5-Ph-IAA, total DNA was extracted, and viral DNA was quantified as described in the figure legend for Fig. 2. **, *p*<0.01. NS, not significant. (B) Relative cell number after treatment with or without chloroquine. The number of cells was manually counted by using a hemocytometer. The number of cells at 0 h was set as 1. NS, not significant. (C and D) Knockdown of cGAS, STING, and autophagy-related molecules rescued the reduction of viral DNA upon LANA depletion. iSLK-OsTIR1-mAID-LANA BAC16 cells were transfected with 25 nM siRNA against indicated molecules for 48 h. The cells were then treated with 2 μM of 5-Ph-IAA for another 24 h, and followed by the indicated analyses. (C) Total DNA was extracted, and viral DNA was quantified as described above. **, *p*<0.01. *, *p*<0.05. (D) Total RNA was extracted, and mRNA expression of target genes was quantified by real-time qPCR. GAPDH was used as an internal standard for normalization. **, *p*<0.01.

cGAS is an invaluable DNA sensor for cellular innate immunity, and plays an important role in the detection of dsDNA derived from pathogens as well as self-origin (42). Upon binding to dsDNA, cGAS produces a second messenger 2’3’-cyclic GMP-AMP (2’3’-cGAMP) that activates the downstream protein, Stimulator of Interferon Genes (STING) localized on the ER membrane. STING then migrates from the ER to the ER-Golgi intermediate compartment (ERGIC) and/or Golgi apparatus where it stimulates the interferon production (53). In addition to interferon production, a recent study revealed that STING induces autophagy (43). We therefore asked if viral DNA is degraded by cGAS-STING-mediated autophagy when LANA is depleted. We transfected siRNA against cGAS, STING, and several autophagy-related genes (e.g., ATG2A) to see if the knocking down of these genes rescues the viral DNA reduction upon LANA depletion. We first confirmed the gene silencing, and the results showed that siRNAs efficiently reduced targeted mRNA expression in iSLK-OsTIR1-mAID-LANA BAC16 cells. The knockdown of cGAS, STING, ATG2A, ATG2B, ATG5, and ATG18 partially but significantly rescued the viral DNA reduction upon LANA depletion (**Fig. 4C, D**).

### cGAS is required for the viral DNA reduction

To further confirm if cGAS is required for viral DNA reduction, we employed a cGAS-null cell line. It is known that human embryonic kidney 293 (HEK293) cell and its sublineages lack the expression of cGAS (54). We first confirmed that iSLK-OsTIR1-mAID-LANA BAC16 cells express cGAS but not in 293FT-OsTIR1-mAID-LANA BAC16 cells (**Fig. 5A**). We next verified if the AID system is functional in 293FT-OsTIR1-mAID-LANA BAC16 cell. The cells were treated with 5-Ph-IAA for 24 h, and lytic reactivation was induced with 20 ng/ml of 12-*O*-Tetradecanoylphorbol-13-acetate (TPA) for another 24 h. As shown in Fig. 5B, LANA was depleted in response to 5-Ph-IAA. Interestingly, LANA depletion again attenuated the expression of lytic proteins, K-Rta and ORF57, similar to what we observed in iSLK-OsTIR1-mAID-LANA BAC16 cell. The results suggested that degradation of episome is not the only reason for the attenuation of lytic gene transcriptions. We then quantified viral DNA copy at 24 h after the addition of 5-Ph-IAA, and the results showed that viral DNA copy was decreased to approximately 63% of that in control cells (**Fig. 5C**). We also counted the number of cells in the same condition, and the results demonstrated that relative cell number of 293FT-OsTIR1-mAID-LANA BAC16 cells at 24 h time point was approximately 1.7 (**Fig. 5D**). If viral DNA does not replicate due to the lack of LANA and is diluted by cell division, the relative viral DNA copy would be the reciprocal of relative cell number. If the relative cell number is 3.0, the theoretical copy number would be 1/3 = 0.33. In the present study, the relative cell number was 1.72, thus the theoretically expected copy number is 1/1.72 = 0.58. The relative episome number (0.63) was slightly larger than 0.58, suggesting that viral episomes were likely to be depleted only by cell division in 293FT-OsTIR1-mAID-LANA BAC16 cells. Taken together, these results suggested that LANA is actively protecting KSHV episome from cGAS-STING axis, probably by the formation of LANA nuclear bodies (**Fig. 5E**).

**Fig. 5.**
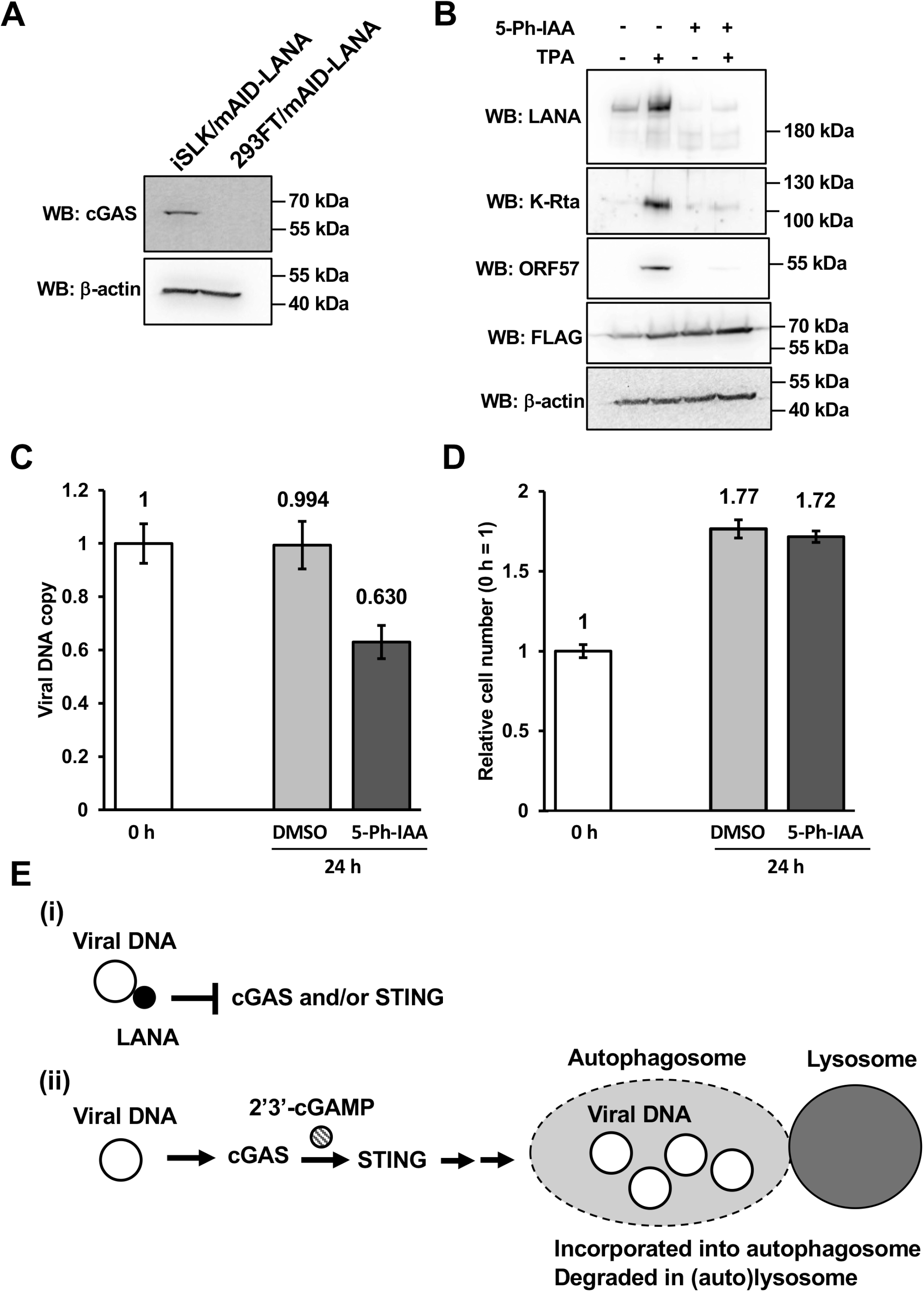
cGAS is required for the viral DNA reduction upon LANA depletion. (A) iSLK-OsTIR1-mAID-LANA BAC16 cells express cGAS but not in 293FT-OsTIR1-mAID-LANA BAC16 cells. Expression of cGAS was assessed by Western blotting. (B) The AID system is functional in 293FT cells. 293FT-OsTIR1-mAID-LANA BAC16 cells were treated with or without 2 μM of 5-Ph-IAA for 24 h, treated with or without 20 ng/ml of 12-*O*-Tetradecanoylphorbol-13-acetate (TPA) for another 24 h, and then the expression of viral proteins was assessed by Western blotting. (C and D) 293FT-OsTIR1-mAID-LANA BAC16 cells were treated with DMSO (0.1%) or 2 μM 5-Ph-IAA for 24 h, and then followed by the indicated analyses. (C) Total DNA was extracted, and viral DNA copy was measured by real-time qPCR. 18S rRNA was used as an internal standard for normalization. (D) Cells were trypsinized and suspended in 1% BSA, 0.1 mM EDTA in PBS. The number of cells was manually counted by using a hemocytometer. The number of cells at 0 h was set as 1. (E) Hypothetical mechanism for how LANA protects viral DNA. (i) When LANA is present, LANA inhibits cGAS and/or STING to protect viral genomic DNA from degradation. (ii) When LANA is absent, cGAS produces 2’3’-cGAMP to stimulate STING. STING then induces autophagy. Viral genomic DNA is then delivered to the lysosome and degraded.

## DISCUSSION

In the present study, we applied the AID system for studying the role of KSHV LANA in latency maintenance. By integrating a mAID-tag into KSHV BAC16 using a recombination-based method, we demonstrated the utility of this approach to study the function of a KSHV viral protein in infected cells. RNA interference (RNAi)-mediated gene silencing has been utilized to study the function of specific genes/proteins of interest in various organisms in the past two decades. Some genes, however, cannot be knocked down because they are essential for cell viability. In addition, the RNAi approach relies on the turnover (half-life) of the target protein for efficient reduction in target protein levels. For silencing of most proteins, RNAi is usually effective only after 48-72 h of siRNA transfection (55). On the other hand, the inducible protein degradation approach allows us to quickly deplete the target protein (37, 38). Indeed, we showed that mAID-LANA was successfully depleted within 1.5 h after the addition of 5-Ph-IAA. We believe that the rapid depletion of target protein enables us to identify the biological effects more directly. After learning that mAID-LANA can be depleted rapidly and efficiently, we tagged several other viral proteins with mAID using the same recombination approach. We found that, however, K-Rta (ORF50) could not be depleted by treatment with 5-Ph-IAA in a similar time frame. In this method, the mAID-tagged target protein must be polyubiquitinated by SCF-OsTIR1 E3 ubiquitin ligase for proteasomal degradation. Accordingly, the abundance of target protein expression and accessibility of the E3 ligase to mAID-tag would have a significant impact on the outcome of target protein degradation. We speculate that having highly concentrated homo-multimers at specific chromosomal loci might be a reason for the great success of LANA degradation.

Previous studies revealed that LANA plays an important role in maintaining latency by repressing the transcriptional activity of the K-Rta promoter (56, 57). K-Rta is a key transcriptional regulator that controls the switch from latency to lytic replication (8). LANA downregulates K-Rta’s promoter activity in transient reporter assays, thus repressing K-Rta-mediated transactivation (58). LANA also physically interacts with K-Rta *in vitro* and in KSHV-infected cells (58). In addition, a study showed that KSHV can undergo spontaneous lytic reactivation, and K-Rta transcription level was increased when LANA expression was knocked down with a specific siRNA (59). These results establish a model in which LANA is actively suppressing viral lytic replication by antagonizing the functions of K-Rta. In the present study, we showed that the depletion of LANA itself did not induce lytic gene expression in iSLK cells. We found that LANA depletion instead further decreased viral lytic gene expression, which is triggered by doxycycline and sodium butyrate treatment. We noticed that KSHV latent chromatin is heavily modified with heterochromatin mark (H3K27me3) in iSLK cells, when we compared with KSHV episomes in primary effusion lymphoma cells. This may partly explain why the depletion of LANA alone did not trigger reactivation. Degradation of KSHV episome demonstrated here also complicated the analyses. Detail time course as well as increasing resolution of studies via isolation from reactivating from the dish would clarify LANA’s role in lytic gene transcription during reactivation.

Viral genomic copy number was reduced to approximately 25% of control within 24 h of 5-Ph-IAA incubation; the results strongly suggest that the KSHV episome is targeted by an active DNA degradation pathway in the absence of LANA. Sensing of viral constituents is the critical step in the host innate immune response against viruses (60). Several innate immune pathways have been identified, including Toll-like receptors (TLRs) (61), pattern recognition receptors (PRRs) (62), and cGMP-AMP synthase (cGAS) (63). Among the molecules involved in innate immunity, cGAS has emerged in recent years as a non-redundant dsDNA sensor important for detecting many pathogens (63). A recent study demonstrated that LANA binds to cGAS, and inhibits the cGAS-STING pathway and thereby antagonizes the cGAS-STING-mediated restriction of KSHV lytic replication (64). In the present study, we found that the viral genomic DNA degradation upon LANA depletion was mediated by the lysosome. In addition, cGAS, STING, and autophagy play a role in viral DNA degradation. Particularly, cGAS is required for the viral genomic DNA degradation in the absence of LANA (a model depicted in **Fig. 5**). A remaining question would be; how LANA antagonizes cGAS to protect viral DNA? One possibility is that LANA binds to cGAS and prevents cGAS to recognize dsDNA. However, this mechanism would require large copies of LANA molecules to absorb cGAS molecules neighboring to KSHV episomes. Locally concentrated LANA via TR binding and formation of nuclear bodies may therefore facilitate the function. Our previous study also suggests that LANA may be involved in the formation of 3D episome structure through tight binding with TRs via a genomic looping (65). We speculate that rapid loss of LANA may therefore disrupt such 3D genomic structure to facilitate the recognition of viral DNA by cGAS. It is very exciting and important to clarify how LANA plays a role in protecting latent viral episomes from degradation. Nonetheless, this study again suggested that LANA is a very attractive target for therapeutic intervention for KSHV-associated diseases.

In summary, we successfully adapted the AID approach to recombinant KSHV studies and demonstrated the utility of the approach to assess viral protein function. Our results suggested that LANA plays an important role in protecting latent viral episome from innate immune DNA sensing.

## MATERIALS AND METHODS

### Reagents

Dulbecco’s modified minimal essential medium (DMEM), fetal bovine serum (FBS), phosphate-buffered saline (PBS), trypsin-EDTA solution, 100x penicillin-streptomycin-L-glutamine solution (Pen-Strep-L-Gln), Alexa 405-conjugated secondary antibody, Alexa 555-conjugated secondary antibody, Alexa 647-conjugated secondary antibody, SlowFade Gold antifade reagent, Lipofectamine 2000 reagent, and high-capacity cDNA reverse transcription kit were purchased from Thermo Fisher (Waltham, MA, USA). Puromycin solution and Zeocin solution were obtained from InvivoGen (San Diego, CA, USA). Hygromycin B solution was purchased from Enzo Life Science (Farmingdale, NY, USA). Anti-FLAG M2 mouse monoclonal and anti-FLAG rabbit polyclonal antibodies, anti-LANA rat monoclonal antibody, anti-β-actin mouse monoclonal antibody, and polyvinylidene difluoride (PVDF) membrane were purchased from Millipore-Sigma (Burlington, MA, USA). Anti-K8α mouse monoclonal and anti-ORF57 mouse monoclonal antibodies were purchased from Santa Cruz Biotechnology (Santa Cruz, CA, USA). Anti-cGAS rabbit monoclonal antibody was purchased from Cell Signaling Technology (Danvers, MA, USA). Anti-mAID mouse monoclonal antibody was purchased from MBL (Tokyo, Japan). Anti-K-Rta rabbit polyclonal antibody was described previously (66). 5-Ph-IAA was purchased from MedChemExpress (Monmouth Junction, NJ, USA). cOmplete protease inhibitor cocktail tablets were purchased from Roche (Basel, Switzerland). ON-TARGETplus siRNAs were purchased from Horizon Discovery Ltd. (Cambridge, UK). The Quick-RNA Miniprep kit was purchased from Zymo Research (Irvine, CA, USA) and QIAamp DNA mini kit was purchased from QIAGEN (Germantown, MD, USA). The NucleoBond Xtra BAC kit was purchased from TaKaRa Bio (Kusatsu, Shiga, Japan). All other chemicals were purchased from Millipore-Sigma or Fisher Scientific unless otherwise stated.

### Cell culture

iSLK cells were maintained in DMEM supplemented with 10% FBS, 1x Pen-Strep-L-Gln, and 2 μg/ml puromycin at 37°C with air containing 5% carbon dioxide. iSLK cell is a derivative of SLK cell which was originally isolated from gingival endothelial tissue, and was transduced with retroviruses expressing rtTA and K-Rta (67). 293FT cells were grown in DMEM supplemented with 10% FBS and 1x Pen-Strep-L-Gln at 37°C with air containing 5% carbon dioxide. iSLK.219 cells were maintained in DMEM supplemented with 10% FBS, 10 μg/ml puromycin, 400 μg/ml hygromycin B, and 250 μg/ml G418. iSLK cells harboring BAC16 WT were maintained in DMEM supplemented with 10% FBS, 1x Pen-Strep-L-Gln, 1,000 μg/ml hygromycin B, and 2 μg/ml puromycin at 37°C with air containing 5% carbon dioxide.

### Construction of mAID-LANA KSHV BAC16

Recombinant KSHV was prepared by following a protocol for *en passant* mutagenesis with a two-step markerless red recombination technique (44). Briefly, mAID-coding sequence was first synthesized (IDT DNA gBlock) and cloned into a pBlueScript SK vector. The sequence is indicated in Table I. The pEPkan-S plasmid (68) was also used as a source of the kanamycin cassette, which includes an I-SceI restriction enzyme site at the 5’ end of the kanamycin resistance gene-coding region. The kanamycin cassette was amplified with primer pairs listed in Table I. An amplified kanamycin cassette was then cloned into the mAID-coding region. The resulting plasmid was used as a template for another round of PCR to prepare a transfer DNA fragment for markerless recombination with BAC16. The PCR fragment was electroporated into E. coli strain GS1783 harboring wild-type BAC16 using Bio-Rad E. coli Pulser. The electroporated E. coli was seeded onto a Luria broth (LB) agar plate containing 30 µg/ml chloramphenicol and 30 µg/ml kanamycin. Positive co-integrates were identified by colony PCR with appropriate primer pairs. Two independent co-integrates were cultured in LB containing 30 µg/ml chloramphenicol and 1% L-arabinose to induce I-SceI, and the E. coli was seeded onto an LB agar plate containing 30 µg/ml chloramphenicol and 1% L-arabinose. Positive clones were identified by colony PCR. The recombination junction and adjacent genomic regions were amplified by PCR, and the resulting PCR fragments were directly sequenced with the same primers to confirm in-frame insertion into the BAC DNA. Two independent BAC clones were generated as biological replicates. BAC DNA was extracted from *E. coli* using the NucleoBond Xtra BAC kit according to the manufacturer’s protocol. iSLK cells and 293FT cells seeded in a 10-cm dish were transfected with approximately 10 μg of purified mAID-LANA BAC16 using Lipofectamine 2000 reagent and selected with 1,000 μg/ml hygromycin B. iSLK cells and 293FT cells harboring mAID-LANA BAC16 were further transduced with recombinant lentivirus expressing FLAG-tagged OsTIR1 F74G protein, and selected with 500 μg/ml zeocin. The resulting cells were maintained in DMEM supplemented with 10% FBS, 1x Pen-Strep-L-Gln, 2 μg/ml puromycin, 1,000 μg/ml hygromycin B, and 100-200 μg/ml zeocin.

### RNA interference (RNAi)

iSLK-OsTIR1-mAID-LANA BAC16 cells were transfected with 25 nM siRNA against indicated gene using Lipofectamine 2000 reagent. The cells were treated with or without 5-Ph-IAA for another 24 h, and then total DNA was extracted as described elsewhere.

### Western blotting

Cells were lysed in SUMO buffer consisting of 50 mM Tris-HCl (pH6.8), 1% SDS, 10% glycerol, and 1x protease inhibitor cocktail. Total cell lysates were boiled in SDS-PAGE sample buffer, subjected to SDS-PAGE, and subsequently transferred onto a PVDF membrane using a wet transfer apparatus (Bio-Rad, Hercules, CA, USA). The final dilutions of the primary antibodies were 1:500 for anti-LANA, 1:1,000 for anti-mAID, 1:2500 for anti-K-Rta, 1:3,000 for anti-β-actin, 1:2,000 anti-FLAG (M2 mouse monoclonal), 1:200 for anti-ORF57, and 1:200 for anti-K8α. Membrane washes and secondary antibody incubations were performed as described previously (45).

### Real-time qPCR

Total RNA was isolated using a Quick-RNA Miniprep kit according to the manufacturer’s protocol. First-strand cDNA was synthesized using a high-capacity cDNA reverse transcription kit according to the manufacturer’s protocol. Gene expression was analyzed by real-time qPCR using specific primers for KSHV ORFs designed by Fakhari and Dittmer (69). We used 18S rRNA as an internal standard to normalize viral gene expression.

### Immunofluorescence microscopy

Cells grown on 22 x 22 mm glass coverslips were fixed with 4% paraformaldehyde in PBS for 20 min, permeabilized with 0.2% Triton X-100 in PBS for 20-30 min, and then blocked with 2% bovine serum albumin in PBS. The fixed cells were incubated with diluted primary antibody solution, followed by diluted secondary antibody solution. The cells on coverslips were then mounted on glass slides using SlowFade reagent and observed using a Keyence BX-Z710 fluorescence microscope (Osaka, Japan). The final dilutions of the primary antibodies were 1:100 for anti-LANA, 1:200 for anti-FLAG (rabbit polyclonal), and 1:100 for K8α.

### Flow cytometry

iSLK cells were infected with purified mAID-LANA virus at a multiplicity of infection (MOI) of 10. At approximately 48 h after infection, the cells were trypsinized, fixed with 2% paraformaldehyde, and suspended in 1% bovine serum albumin in PBS. Samples were acquired on Accuri C6 flow cytometer (BD Bioscience, Franklin Lakes, NJ, USA) and analyzed using FlowJo software (BD Bioscience) as described previously (45).

### Quantification of viral genomic DNA copy in cells

iSLK-OsTIR1-mAID-LANA BAC16 cells were seeded into 6-well plates and treated with 2 μM 5-Ph-IAA or 0.1% dimethyl sulfoxide (DMSO) for 24 h. The cells were trypsinized and suspended in 200 μl of PBS. Total DNA was extracted using a QIAamp DNA mini kit according to the manufacturer’s protocol. Five μl of eluate was used for real-time qPCR to determine viral genomic DNA copy number. We used 18S rRNA as an internal standard to normalize viral genome copy numbers.

### Quantification of progeny virus production

iSLK-OsTIR1-mAID-LANA BAC16 cells were seeded into 6-well plates, and treated with 2 μM 5-Ph-IAA or 0.1% dimethyl sulfoxide (DMSO) for 24 h. The cells were then treated with 1 μg/ml of doxycycline and 1.5 mM sodium butyrate for another 96 h together with 5-Ph-IAA. Two hundred μl of cell culture supernatant was treated with 12 μg/ml DNase I for 15 min at room temperature to degrade DNAs that were not correctly encapsidated. The reaction was stopped by the addition of 5 mM EDTA, followed by heating at 70°C for 15 min. Viral DNA was then purified using a QIAamp DNA Mini kit according to the manufacturer’s protocol. Five μl of eluate was used for real-time qPCR to determine viral genomic DNA copy number.

### Statistical analysis

Results are shown as means ± S.E.M. from at least three independent experiments. Data were analyzed using an unpaired Student’s *t*-test. An FDR-corrected *p-*value less than 0.05 was considered statistically significant.

## ACKNOWLEDGEMENTS

This research was supported by Public Health Service grants from the National Cancer Institute (CA225266, CA232845) to Y.I.

